# A distributed brain network predicts general intelligence from resting-state human neuroimaging data

**DOI:** 10.1101/257865

**Authors:** Julien Dubois, Paola Galdi, Lynn K. Paul, Ralph Adolphs

## Abstract

Individual people differ in their ability to reason, solve problems, think abstractly, plan and learn. A reliable measure of this general ability, also known as intelligence, can be derived from scores across a diverse set of cognitive tasks. There is great interest in understanding the neural underpinnings of individual differences in intelligence, since it is the single best predictor of longterm life success. The most replicated neural correlate of human intelligence to date is total brain volume; however, this coarse morphometric correlate says little about function. Here we ask whether measurements of the activity of the resting brain (resting-state fMRI) might also carry information about intelligence. We used the final release of the Young Adult Human Connectome Project (N=884 subjects after exclusions), providing a full hour of resting-state fMRI per subject; controlled for gender, age, and brain volume; and derived a reliable estimate of general intelligence from scores on multiple cognitive tasks. Using a cross-validated predictive framework, we predicted 20% of the variance in general intelligence in the sampled population from their resting-state connectivity matrices. Interestingly, no single anatomical structure or network was responsible or necessary for this prediction, which instead relied on redundant information distributed across the brain.

## Introduction

Most psychologists agree that there is, in addition to specific cognitive abilities, a very general mental capability to reason, think abstractly, solve problems, plan, and learn across domains [1]. This ability, intelligence, does not refer to a person’s sheer amount of knowledge but rather to their ability to recognize, acquire, organize, update, select and apply this knowledge [2], to reason and make comparisons [3]. There are large and reliable individual differences in intelligence across species: some people are smarter than others, and some rats are smarter than others [4]. What’s more, these differences matter. Intelligence is one of the most robust predictors of conventional measures of educational achievement [5], job performance [6], socioeconomic success [7], social mobility [8], health [9] and longevity [10,11]; and as life becomes increasingly complex, intelligence may play an ever increased role in life outcome [2]. Despite this overwhelming convergence of evidence for the construct of “intelligence” there is considerable debate about what it really is, how best to measure it, and in particular what predictors and mechanisms for its variability across individuals we could find in the human brain.

### Measuring intelligence: the structure of cognitive abilities

Intelligence tests are among the most reliable, and valid, of all psychological tests and assessments [1]. Psychologists are so confident in the psychometric properties of intelligence tests that, almost 100 years ago, Edwin G. Boring famously wrote: “Intelligence is what the tests test.” [12]. A comprehensive modern intelligence assessment (such as the Wechsler Adult Intelligence Scale, Fourth Edition or WAIS-IV [13]) comprises tasks that assess several aspects of intelligence: some assess verbal comprehension (e.g. word definition, general knowledge, verbal reasoning), some assess visuo-spatial reasoning (e.g., puzzle construction, matrix reasoning, visual perception), some assess working memory (e.g., digit span, mental arithmetic, mental manipulation), and some assess mental processing speed (e.g., reaction time for detection). The scores on all of these tasks are tallied and compared to a normative, age-matched sample to calculate a standardized Full Scale Intelligence Quotient (FSIQ) score.

One of the most important findings in intelligence research is that performances on all these seemingly disparate tasks -- and many other cognitive tasks -- are positively correlated: individuals who perform above average on, say, visual perception, also tend to perform above average on, say, word definition. Spearman [14] described this phenomenon as the *positive manifold*, and since then it has been described in a number of non-human animal species [15]. To account for this empirical observation, he posited the existence of a general factor of intelligence, the ‘g-factor’, or simply ‘*g*’, which no single task can perfectly measure, but which can be derived from performance on several cognitive tasks through factor analysis. The g-factor captures around half of people’s intellectual differences [16] and shows good reliability across sets of cognitive tasks [17]. The empirical observation of the positive manifold is well-established [18,19]). The descriptive value of Spearman’s *g* is beyond doubt, however its interpretation -- that psychometric *g* reflects a general aspect of brain functioning -- was challenged early on [20] and remains a topic of debate to this day among intelligence researchers [21–23]. The leading alternative theory posits that each cognitive test involves several mental processes, and that the sampled mental processes overlap across tests; in this situation, performance on all tests appears to be positively correlated (“Process Overlap Theory” [19]). The common factor, in this framework, is a consequence of the positive manifold, rather than its cause. There is unfortunately no statistical means of distinguishing between alternative theories on the basis of the psychometric data alone [22].

### The search for biological substrates of intelligence

Differential psychology -- the psychological discipline that studies individual differences between people and, increasingly, between individuals in non-human animal species (as illustrated in this journal issue) -- has three main aims: to describe the trait of interest accurately, to establish its impact in real life, and to understand its ætiology, including its biological basis [24]. Much headway has been made with respect to the first two aims for intelligence in humans, as described above. The third aim, despite much effort, has remained elusive.

Individual differences in intelligence are relatively stable: one of the best predictors of intelligence in old age is - perhaps unsurprisingly - intelligence in childhood [25]. Intelligence has a strong genetic component [5,26,27]; Genome-Wide Association Studies (GWAS) suggest that intelligence is highly polygenic (no single gene accounts for a large fraction of the variance) [28,29]. While high heritability points to genes as a biological substrate of intelligence differences, individual differences in cognitive ability are instantiated in brain function. Arguably, studying this measure -- the most proximal substrate of intelligent behavior -- may yield more direct insight about the actual mechanisms of intelligence than genetic studies have so far. Since intelligence is a relatively stable trait, its aetiology should be found in stable aspects of brain function, and hence also in aspects of brain structure.

Most of the data on the neural basis of intelligence in humans comes from neuroimaging and from lesion studies. Kievit et al. [30] recently described the current state of the neuroscience of intelligence as “an embarrassment of riches” -- a plethora of neuroimaging-derived properties of the brain, structural and functional, have been linked to intelligence over the years, albeit with largely dubious reliability and reproducibility (see below). It is worth noting that the naive search for a simple neurobiological correlate of intelligence faces a major theoretical hurdle, which is best understood through analogy [30]. Imagine a researcher trying to find the biological basis of the construct of “physical fitness”. If they search for a single physical property, they would likely fall short of their goal. Indeed, physical fitness is a composite of several physical properties (such as cardiorespiratory endurance, muscular strength, muscular endurance, body composition, flexibility), and cannot be equated with any single one, or even any specific combination of these factors. It is very likely that intelligence is of a similar composite nature, as suggested by genetic data [28]. Furthermore, individuals may score identically on an IQ test by using different cognitive strategies, or different brain structures [24]. This is an important picture to hold in mind as one searches for neural correlates of intelligence.

There are several in-depth reviews of the neurobiological substrates of intelligence, to which the interested reader is referred for a complete treatment, including structural studies [24,31–33]. Extant functional neuroimaging studies (using EEG, PET, rCBF and fMRI) have been summarized as supporting the notion that intelligence is a network property of the brain, related to neural efficiency [31,34,35]. Foundational studies had very low sample sizes; more recent, better powered studies have found correlations between intelligence and the global connectivity of a small region in lateral prefrontal cortex (N=78) [36]; the nodal efficiency of hub regions in the salience network (N=54) [37]; the modularity of frontal and parietal networks (N=309) [38]; and several other somewhat disparate reports.

### The current study

Extant literature on functional MRI-based correlates of intelligence suffers from the same caveats as most fMRI-based individual differences research to date [39]: small sample sizes, and lack of out-of-sample generalization. A predictive framework was first used in a recent study [40], which found that fluid intelligence as estimated from a short version of Raven’s Progressive Matrices could be predicted from functional connectivity matrices in an early, relatively small-sample release (N=118) of the Human Connectome Project (HCP) dataset [41], with a correlation r_LOSO_=0.50 between observed and predicted scores (the LOSO subscript denotes the use of a leave-one-subject-out cross-validation framework). Since this early report, the authors revised the effect size down to r_LOSO_=0.22 using later data releases (N=606) [42]. Recent work in our group, in which we further control for confounding effects of age, gender, brain size and motion, as well as use a leave-one-family-out cross-validation framework (LOFO) instead of the original leave-one-subject-out framework (thus accounting for the family structure of the HCP dataset, see **Methods**), further revised the effect size down to about r_LOSO_=0.09 using methods matched as closely as possible to the original study [40]; yet, using improvements including better inter-subject alignment and multivariate modeling, we found r_LOSO_=0.263 (N=884) [43]. Note that this effect size is comparable to recent estimates of the relationship between brain size and intelligence [44]. Though the explained variance is small, it would in fact fall around the 65^th^ percentile of correlations observed in individual differences research [45] (with the caveat that r_LOFO_ correlation is derived from a cross-validation procedure, which breaks the assumption of independence between individual data points).

According to recent guidelines [46], the assessment of general intelligence with HCP’s 24-item Progressive Matrices would be considered of “fair” quality (1 test, 1 cognitive dimension, testing time less than 19 minutes), and would be expected to correlate with general intelligence in the range 0.50 to 0.71. This rather low measurement quality itself is likely to attenuate the magnitude of the relationship between neural data and general intelligence. Fortunately, there are several other measures of cognition in the HCP, which we here decided to leverage to derive a better estimate of general intelligence -- one that would meet criteria for an “excellent” quality measurement (more than 9 tests, more than 3 dimensions, testing time more than 40 minutes), and thus be expected to correlate with the general factor of intelligence above 0.95. Our main aims in this study were to: i) predict an excellent estimate of general intelligence from resting-state functional connectivity in a large sample of subjects from the HCP; ii) depending on the success of i), gain some anatomical insight on which functional connections matter for these predictions. The current study paves the way for a reliable neuroimaging-based science of intelligence differences (large sample size; predictive framework; valid, reliable psychometric construct).

## Methods

Many of the methods, in particular the preprocessing of fMRI data and the predictive analyses, were developed and described in more detail in our recent publication on personality [43].

### Dataset

We used data from a public repository, the 1200 subjects release of the Human Connectome Project (HCP) [41]. The HCP provides MRI data and extensive behavioral assessment from almost 1200 subjects. Acquisition parameters and “minimal” preprocessing of the resting-state fMRI data is described in the original publication [47]. Briefly, each subject underwent two sessions of resting-state fMRI on separate days, each session with two separate 14 min 24 s acquisitions generating 1200 volumes (customized Siemens Skyra 3 Tesla MRI scanner, TR = 720 ms, TE = 33 ms, flip angle= 52°, voxel size = 2 mm isotropic, 72 slices, matrix = 104 × 90, FOV = 208 mm x 180 mm, multiband acceleration factor = 8). The two runs acquired on the same day differed in the phase encoding direction, left-right and right-left (which leads to differential signal intensity especially in ventral temporal and frontal structures). The HCP data was downloaded in its minimally preprocessed form, i.e. after motion correction, B0 distortion correction, coregistration to T_1_-weighted images and normalization to MNI space.

### Cognitive ability tasks

Previous studies of the neural correlates of intelligence in the HCP [40,42] have relied on the number of correct responses on form A of the 24(+3 bonus)-item Penn Matrix Reasoning Test (PMAT), a test of non-verbal reasoning ability which can be administered in under 10 minutes (mean=4.6, std= 3 min; [48]), and is included in the University of Pennsylvania Computerized Neurocognitive Battery (Penn CNB, [49–51]). The PMAT [52,53] is designed to parallel many of the psychometric properties of the Raven’s Standard Progressive Matrices test (RSPM, originally published in 1938, which comprises 60 items [54]), while limiting learning effects and expanding the representation of the abstract reasoning construct (Ruben Gur, personal communication).

Assessment of cognitive ability in the HCP [55] also includes several tasks from the Blueprint-funded NIH Toolbox for Assessment of Neurological and Behavioral function (http://www.nihtoolbox.org), as well as other tasks from the Penn computerized neurocognitive battery [50]. These other tasks can be leveraged to derive a better measure of the general intelligence factor [46]. We included all cognitive tasks listed in the HCP Data Dictionary, except for: i) the delay discounting task, which is not a measure of ability (i.e. there is not a correct response); and ii) the Short Penn Continuous Performance Test which is about sustained attention rather than cognitive ability, and whose distribution departed too much from normality (data not shown). Our initial selection thus consisted of 10 tasks (NIH Toolbox: dimensional change card sort; flanker inhibitory control and attention; list sorting working memory; picture sequence memory; picture vocabulary; pattern comparison processing speed; oral reading recognition; Penn CNB: Penn progressive matrices; Penn word memory test; variable short Penn line orientation), which are also listed in **Supplementary Table 1** along with a brief description (the descriptions are copied almost word for word from the HCP Data Dictionary). When several outcome measures were available for a given task, we selected the one that best captured ability; when both age-adjusted and unadjusted scores were available, we included the unadjusted scores. Though some of the NIH toolbox scores combine accuracy and reaction time, we only considered accuracies for the Penn CNB tasks (to avoid confounding ability and processing speed; but, see [51]).

### Subject selection

The total number of subjects in the 1200-subject release of the HCP dataset is N=1206. We applied the following criteria to include/exclude subjects from our analyses (listing in parentheses the HCP database field codes). i) Complete neuropsychological datasets. Subjects must have completed all relevant neuropsychological testing (PMAT_Compl=True, NEO-FFI_Compl=True, Non-TB_Compl=True, VisProc_Compl=True, SCPT_Compl=True, IWRD_Compl=True, VSPLOT_Compl=True) and the Mini Mental Status Exam (MMSE_Compl=True). Any subjects with missing values in any of the tests or test items were discarded. This left us with N = 1183 subjects. ii) Cognitive compromise. We excluded subjects with a score of 26 or below on the MMSE, which could indicate marked cognitive impairment in this highly educated sample of adults under age 40 [56]. This left us with N = 1181 subjects (638 females, 28.8 +/− 3.7 y.o., range 22-37 y.o). This is the sample of subjects available for factor analyses. Furthermore, iii) subjects must have completed all resting-state fMRI scans (3T_RS-fMRI_PctCompl=100), which leaves us with N = 988 subjects. Finally, iv) we further excluded subjects with a root-mean-squared frame-to-frame head motion estimate (Movement_Relative_RMS.txt) exceeding 0.15mm in any of the 4 resting-state runs (threshold similar to [40]). This left us with the final sample of N = 884 subjects (475 females, 28.6 +/− 3.7 y.o., range 22-36 y.o.) for predictive analyses based on resting-state data.

### Deriving the general factor of intelligence, *g*

There are several methods in the literature to derive a general factor of intelligence from scores on a set of cognitive tasks. The simplest consists in using a standardized sum score composite; this is the conventional approach when all scores come from a well-validated battery. However, since we are here including scores from two different cognitive batteries (NIH toolbox and Penn CNB), we sought to characterize the structure of cognitive abilities in our sample using factor analysis, and then derive scores for the general factor. We conducted an exploratory factor analysis (EFA), specifying the bi-factor model of intelligence -- a common factor g which loads on all test scores, and several group factors that each load on subsets of the test scores; all latent factors are orthogonal to one another -- using the *psych* (v1.7.8) package [57] in *R* (v3.4.2). We specifically used the *omega* function, which conducts a factor analysis (with maximum likelihood estimation) of the data set, rotates the factors obliquely (using “oblimin” rotation), factors the resulting correlation matrix, then does a Schmid-Leiman transformation [58] to find general factor loadings. Model fit was assessed using several commonly used statistics in factor analysis [59]: the Comparative Fit Index (CFI; should be as close to 1 as possible; values >0.95 are considered a good fit); the Root Mean Squared Error of Approximation (RMSEA; should be as close to 0 as possible; values <0.06 are considered a good fit); the Standardized Root Mean Squared Residual (SRMR; should be as close to 0 as possible; values <0.08 are considered a good fit); and the Bayesian Information Criterion (BIC; better models have lower values, can be negative). Factor scores can be derived using different methods [60], for example the regression method. This approach mimics the one taken by [61]. It is however usually preferable to derive scores from a confirmatory factor analysis (CFA). The main difference between EFA and CFA is that in EFA, observed task scores are allowed to cross-load freely on several group factors, while in CFA such cross-loadings can be forbidden. For the purpose of deriving the general factor of intelligence, there is little difference between using CFA and EFA in practice; we conducted a CFA using the *lavaan* (v0.5-23.1097) package [62] in *R* to verify this (see **Supplementary Figure 1**).

### Assessment and removal of potential confounds

We computed the correlation of the general factor of intelligence *g* with Gender (HCP variable: *Gender*), Handedness and Age (restricted HCP variables: *Handedness, Age_in_Yrs*). We also looked for differences in *g* in our subject sample with variables that are likely to affect FC matrices, such as brain size (we used *FS_BrainSeg_Vol*), motion (we computed the sum of framewise displacement in each run), and the multiband reconstruction algorithm which changed in the third quarter of HCP data collection (*fMRI_3T_ReconVrs*). We then used multiple linear regression to regress these variables from *g* scores and remove their confounding effects.

Note that we did not control for differences in cortical thickness and other morphometric features, which have been reported to be correlated with intelligence (e.g. [63]). These likely interact with FC measures and should eventually be accounted for in a full model, yet this was deemed outside the scope of the present study.

Note also that we also did not consider Educational Achievement (EA) and Socio-Economic Status (SES) as confounds in this analysis, as they have been described as consequences of intelligence and thus controlling for them would remove meaningful variance. A full model considering these variables is a future direction for this work.

### Data preprocessing

We recently explored the effects of several preprocessing pipelines on the prediction of personality factors and PMAT scores in the HCP dataset [43]. Here we adopted the preprocessing pipeline which was found to produce the highest prediction scores in that study. The pipeline reproduces as closely as possible the strategy described in [40] and consists of seven consecutive steps: 1) the signal at each voxel is z-score normalized; 2) using tissue masks, temporal drifts from cerebrospinal fluid (CSF) and white matter (WM) are removed with third-degree Legendre polynomial regressors; 3) the mean signals of CSF and WM are computed and regressed from gray matter voxels; 4) translational and rotational realignment parameters and their temporal derivatives are used as explanatory variables in motion regression; 5) signals are low-pass filtered with a Gaussian kernel with a standard deviation of 1 TR, i.e. 720ms in the HCP dataset; 6) the temporal drift from gray matter signal is removed using a third-degree Legendre polynomial regressor; 7) global signal regression is performed. These operations were performed using an in-house, Python (v2.7.14)-based pipeline (mostly based on open source libraries and frameworks for scientific computing, including SciPy (v0.19.0), Numpy (v1.11.3), NiLearn (v0.2.6), NiBabel (v2.1.0), Scikit-learn (v0.18.1) [64–68]).

### Inter-subject alignment, parcellation, and functional connectivity matrix generation

We used surface-based multi-modally aligned cortical data (MSM-All [69]), together with a parcellation that was derived from this data using an objective semi-automated neuroanatomical approach [70]. The parcellation has 360 nodes, 180 for each hemisphere. These nodes can be attributed to the major resting state networks [71] (**Figure 4a**). Time series extraction simply consisted in averaging data from vertices within each parcel, and matrix generation in pairwise correlating parcel time series (Pearson correlation coefficient). We concatenated time series across runs to derive average FC matrices (REST1: from concatenated REST1_LR and REST1_RL time series; REST2: from concatenated REST2_LR and REST2_RL time series; REST12: from concatenated REST1_LR, REST1_RL, REST2_LR and REST2_RL time series). There are (360 * 359)/2 = 64620 undirected edges in a network of 360 nodes. This is the dimensionality of the feature space for prediction.

### Prediction model

We used a univariate feature filtering approach to reduce the number of features, discarding edges for which the p-value of the correlation with the behavioral score is greater than a set threshold, for example p<0.01 (as in [40]). We then used Elastic Net regression to learn the relationship with behavior; based on our previous work [43] and on the fact that it is unlikely that just a few edges contribute to prediction, we fixed the L1 ratio (which weights the L1- and L2-regularizations) to 0.01, which amounts to almost pure ridge regression. We used 3-fold nested cross-validation (with balanced “classes”, based on a partitioning of the training data into quartiles) to choose the alpha parameter (among 50 possible values) that weighs the penalty term.

### Cross-validation scheme

In the HCP dataset, several subjects are genetically related (in our final subject sample there were 410 unique families). To avoid biasing the results due to this family structure (e.g., perhaps having a sibling in the training set would facilitate prediction for a test subject, if both intelligence and functional connectivity are heritable), we implemented a leave-one-family-out crossvalidation scheme for all predictive analyses.

### Statistical assessment of predictions

Several measures can be used to assess the quality of predictions, which we described in more detail in our previous publication [43]. Here we report the Pearson correlation coefficient between observed scores and predicted scores, the coefficient of determination R^2^, and the related normalized Root Mean Square Deviation (nRMSD).

In a cross-validation scheme, the folds are not independent of each other. This means that statistical assessment of the cross-validated performance using parametric statistical tests is problematic [43,72,73]. Proper statistical assessment should thus be done using permutation testing on the actual data. To establish the empirical distribution of chance, we ran our predictive analysis using 1000 random permutations of the scores (shuffling scores randomly between subjects, keeping everything else exactly the same, including the family structure).

## Results

### A general factor, *g*, accounts for 58% of the covariance structure of cognitive tasks in the HCP sample

All selected cognitive task scores (**Supplementary Table 1**) correlated positively with one another, as expected from the well-known positive manifold (**Figure 1a**). A parallel analysis suggested an underlying 4-factor structure (**Figure 1b**). An exploratory bifactor analysis with a general factor *g* and 4 group factors fit the data very well (CFI=0.990; RMSEA=0.0311; SRMR=0.0201; BIC=-0.519), much better than a single factor model (CFI=0.719; RMSEA=0.1398; SRMR=0.0887; BIC=591.172). The solution is depicted in **Figure 1c**. The four factors can naturally be interpreted as: 1) Crystallized Ability [cry] (PicVocab_Unadj + ReadEng_Unadj); 2) Processing Speed [spd] (CardSort_Unadj + Flanker_Unadj + ProcSpeed_Unadj); 3) Visuospatial Ability [vis] (PMAT24_A_CR + VSPLOT_TC); and 4) Memory [mem] (IWRD_TOT + PicSeq_Unadj + ListSort_Unadj).

**Figure 1.**
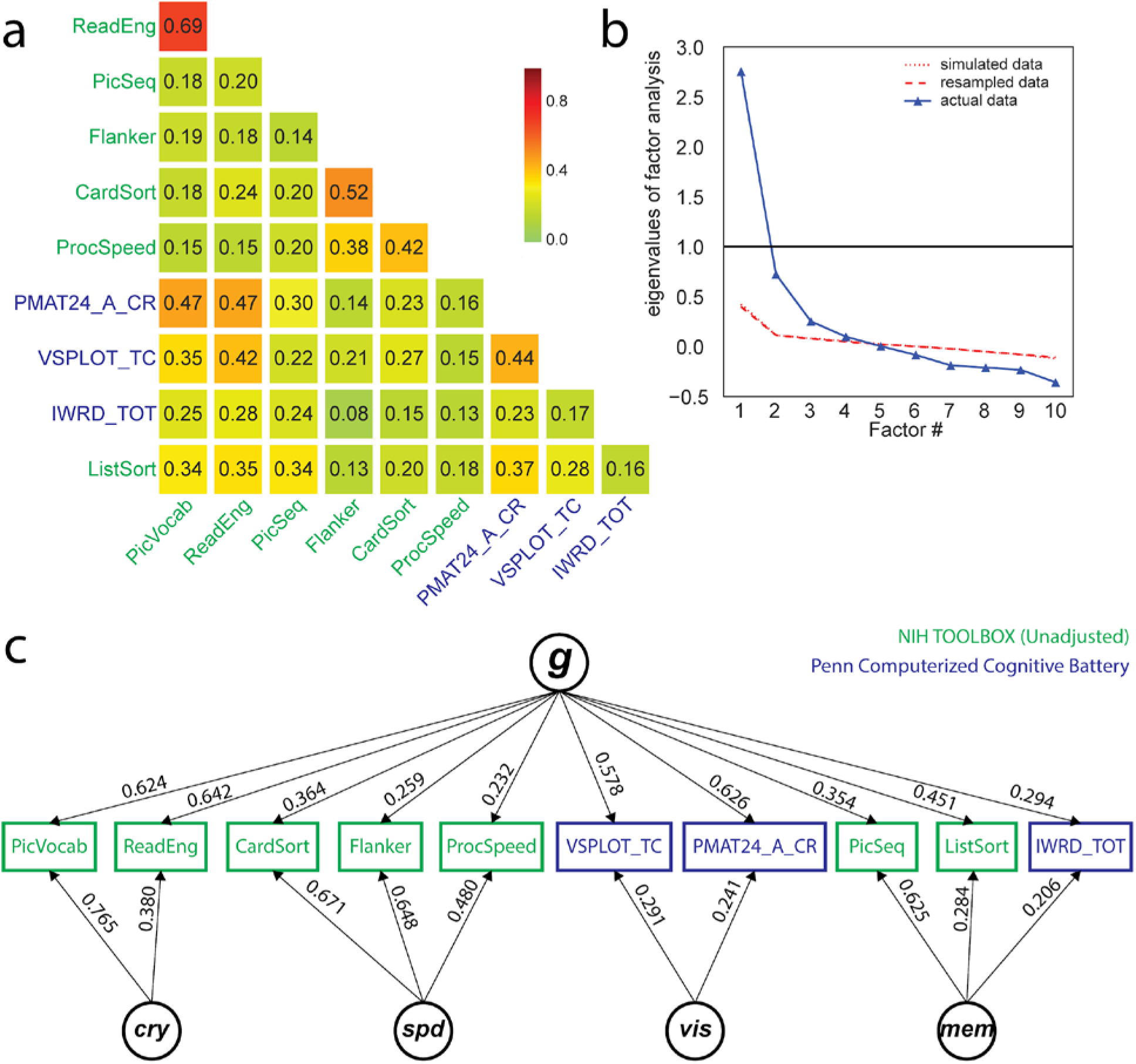
Exploratory factor analysis of select cognitive tasks (Supplementary Table 1) in the Human Connectome Project (HCP) dataset, using N=1181 subjects. **a)** All cognitive task scores correlated positively with one another, reflecting the well-established positive manifold (see also **Supplementary Figure 2** for scatter plots). **b)** A parallel analysis suggested the presence of 4 latent factors from the covariance structure of cognitive task scores. Note that the simulated and resampled data lines are nearly indistinguishable. **c)** A bifactor analysis fit the data well (see fit statistics in text), and yielded a theoretically plausible solution with a general factor (**g**) and four group factors which can be interpreted as crystallized ability (***cry***), processing speed (***spd***), visuospatial ability (***vis***) and memory (***mem***). Loadings less than 0.2 are not displayed.

Across all cognitive task scores, this general intelligence factor accounted for 58.5% of the variance (coefficient omega_hierarchical ω_h [74–76]_), while group factors accounted for 18.2% of the variance (with 15.5% of the variance unaccounted for). Another important metric is coefficient omega subscale ω_s_ [77] which quantifies the reliable variance across the tasks accounted for by each subscale, beyond that accounted for by the general factor; we found ω_s_^cry^=38.7%, ω_s_^spd^=57.4%, ω_s_^vis^=9.9% and ω_s_^mem^=27.8%. While some of these subscale factors account for a substantial proportion of the variance across their respective tasks, their measurement quality is at most fair [46] due to the limited number of constituent tasks; thus we chose to focus on the general factor *g* only in the present study.

Factor scores are indeterminate, and several alternate methods exist to derive them from a structural model [60]. To avoid this issue, most researchers prefer to remain in latent space for further analyses [30]. However, for subsequent analyses we required factor scores for *g*, which we derived using regression-based weights (“Thurstone” method). We compared the general factor scores derived from this exploratory factor analysis (EFA) with a simple composite score consisting of the sum of standardized observed test scores. As expected, we found that the simple composite score correlates highly with the EFA-derived *g* (r=0.91).

### Brain size, gender, and motion are correlated with g

There are known effects of gender [78,79], age [80,81], in-scanner motion [82–84] and brain size [85] on the functional connectivity patterns measured in the resting-state with fMRI. It is thus necessary to control for these variables [86]: indeed, if intelligence is correlated with gender, one would be able to predict some of the variance in intelligence solely from functional connections that are related to gender. The easiest way to control for these confounds is to remove any relationship between the confounding variables and the score of interest in our sample of subjects, which can be done using multiple regression. Note that this approach may be too conservative, and that more work remains to be done on dealing with such confounds (see **Discussion**).

We characterized the relationship between intelligence and each of the confounding variables listed above in our subject sample (**Supplementary Figure 3**). Intelligence was correlated with gender (men scored higher in our sample), age (younger scored higher in our sample -- note the limited age range 22-36 y.o.), and brain size (larger brains scored higher) [44,46]. There was no relationship between handedness and intelligence in our sample (r=2×10^−6^). Motion, quantified as the sum of frame-to-frame displacement over the course of a run (and averaged separately for REST1 and REST2) was correlated with intelligence: subjects scoring lower on intelligence moved more during the resting-state. Note that motion in REST1 was highly correlated (r=0.72) with motion in REST2, indicating that motion itself may be a stable trait, and correlated with other traits. While the interpretation of these complex relationships would require further work outside the scope of this study (using partial correlations, and mediation models, to disentangle effects), it is critical to remove shared variance between intelligence and the primary confounding variables before proceeding further. This ensures that our model is trained specifically to predict intelligence, rather than confounds that covary with it in our subject sample [86]. However there are several other variables that we do not explicitly account for here, for example the Openness personality trait which we previously found to be correlated with intelligence [43].

Another possible confound, specific to the HCP dataset, is a difference in the image reconstruction algorithm between subjects collected prior to and after April 2013. The reconstruction version leaves a notable signature on the data that can make a large difference in the final analyses produced [87]. We found a small, but significant correlation with intelligence in our sample (indicating that subjects imaged with the old reconstruction version were, on average, less intelligent than the ones imaged with the newer reconstruction version). This confound is, of course, a simple sampling bias artifact with no meaning. Yet, this significant artifactual correlation must be removed, by including the reconstruction factor as a confound variable.

Importantly, the multiple linear regression used for removing the variance shared with confounds was fitted on the training data (in each cross-validation fold during the prediction analysis), and then the fitted weights were applied to remove the effects of confounds in both the training and test data. This is critical to avoid any leakage of information, however negligible, from the test data into the training data.

### Resting-state FC predicts 20% of the variance in *g* across subjects

We computed a resting-state functional connectivity matrix for each subject from close to 1 hour of resting-state data (REST12), yielding a very reliable estimate of the stable functional network of each individual -- with 1 h of scan time, a recent study found the test-retest reliability of an individual’s FC matrix to be above r=.96 (see Figure 4c in [88]).

We used a leave-one-family-out cross-validation scheme to train a regularized linear model and predict general intelligence from functional connectivity matrices (features are the 64620 undirected edges), in our sample of 884 subjects. We found a significant correlation between observed and predicted *g* scores (r=0.457, p_1000_<0.001, based on 1000 permutations), a coefficient of determination that differs significantly from chance (R^2^=0.206; p_1000_<0.001), and a normalized root mean square deviation that is significantly lower than its null distribution (nRMSD=0.892;p_1000_<0.001) (**Figure 2**).

**Figure 2.**
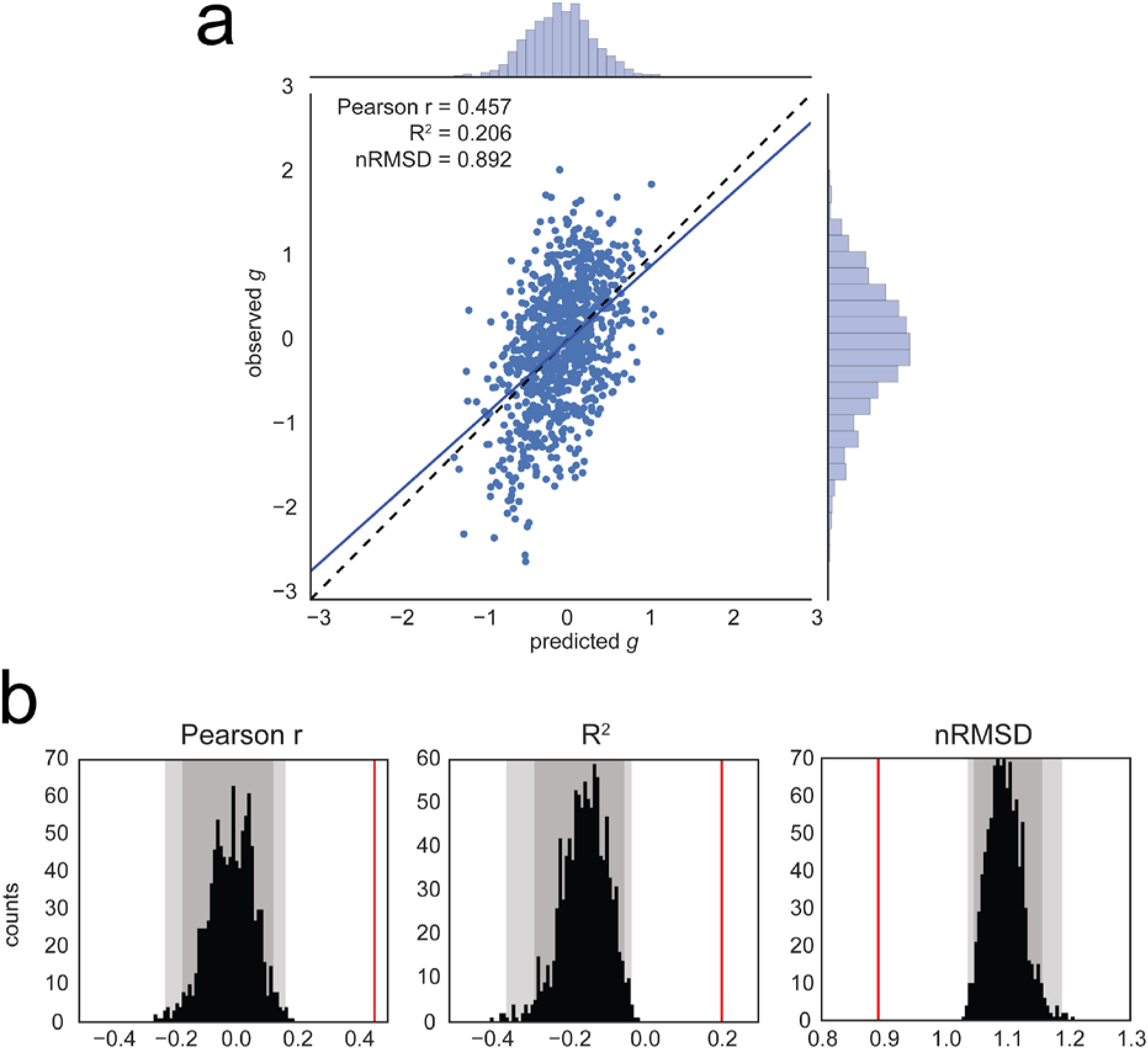
Prediction of the general factor of intelligence *g* from resting-state functional connectivity, averaging all resting-state runs for each subject (REST12, totaling almost 1h of fMRI data). **a)** Observed vs. predicted values of the general factor of intelligence. The regression line had a slope close to 1, as expected theoretically [89]. The correlation coefficient was r=0.457 (REST1 only, r=0.419; REST2 only, r=0.312). **b)** Evaluation of prediction performance according to several statistics and their distributions under the null hypothesis, as simulated through permutation testing (with 1000 surrogate datasets). All fit statistics (red lines) fell far out of the confidence intervals under the null hypothesis. *Left:* the correlation between observed and predicted values; *Center:* the coefficient of determination R^2^, interpretable as the proportion of explained variance; *Right:* the normalized Root Mean Square Deviation nRMSD, which indicates the average difference between observed and predicted scores. Faint gray shade: p<0.05; darker gray shade: p<0.01, permutation tests.

For comparison, we previously found that the prediction of intelligence as estimated by PMAT24_A_CR scores [43] captured less variance (r=0.263,p_1000_<0.001; R^2^=0.047, p_1000_<0.001; nRMSD=0.977, p_1000_<0.001). It is likely that the moderating effect of inferior measurement quality with the PMAT24_A_CR as compared to our factor-derived *g* limited prediction performance [46].

Similarly, using only 30min of resting-state data (one session, two runs) to derive functional connectivity matrices had a moderating effect on prediction performance. With 30 minutes of scan time, test-retest reliability of FC matrices falls to about r=.92 (according to Figure 4c in [88]). Predicting *g* using REST1, we found r=0.419, R^2^=0.170, nRMSD=0.912; using REST2, we found r=0.312, R^2^=0.067, nRMSD=0.966.

### Predictive edges are distributed in FPN, CON, DMN, and VIS networks

We have demonstrated that a substantial and statistically highly significant amount of variance (about 20%) in general intelligence *g* across our sample of subjects can be predicted from resting-state functional connectivity. Is there a specific set of edges (connectivity between specific anatomical parcels) in the brain that carry most of the information? The Parieto-Frontal Integration Theory (P-FIT) of intelligence [31] postulates roles for cortical regions in the prefrontal (Brodmann areas (BA) 6, 9–10, 45–47), parietal (BA 7, 39–40), occipital (BA 18–19), and temporal association cortex (BA 21, 37).

To address this question, we used a descriptive network selection/elimination approach. We focused on the 7 major resting-state networks [90]; Ito and colleagues recently assigned each region of the parcellation used here [70] to these functional networks using the Generalized Louvain method for community detection with resting-state fMRI data [71] (**Figure 3a**). We verified that this published network assignment indeed clustered regions that have similar connectivity patterns at the level of single subjects (**Figure 3b**). We then asked how well we could predict *g* keeping edges within only one network 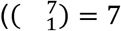 combinations, **Figure 3c**), or within/between two networks 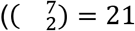combinations, **Figure 3d**). For all these analyses we carried out exactly the same methods as described above (including the univariate feature selection step), but training and testing on a reduced feature space (selecting edges according to networks). Prediction performance within a single network, or with two networks, was much lower compared to performance with the full set of edges (one network, maximum performance r=0.327; two networks, maximum performance r=0.373); however, some networks were found to carry more information than others: the most informative networks were CON, DMN, FPN and VIS, while DAN, AUD and SMN carried very little information. These results are in good agreement with the P-FIT [31]; in particular, in addition to the eponymous frontal and parietal regions which have been reported in other studies already [40,61], we evidenced information in the VIS network as postulated by P-FIT. We next explored how the removal of networks affected prediction: we “lesioned” a single network 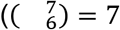, **Figure 3f**) or two networks (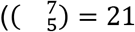, **Figure 3e**). We found that lesioning one or two networks had a very small effect on the prediction of *g* (lesion one network, minimum performance r=0.409; lesion two networks, minimum performance r=0.373), indicating there is distributed and redundant information about *g* in functional connectivity patterns across several brain networks.

**Figure 3.**
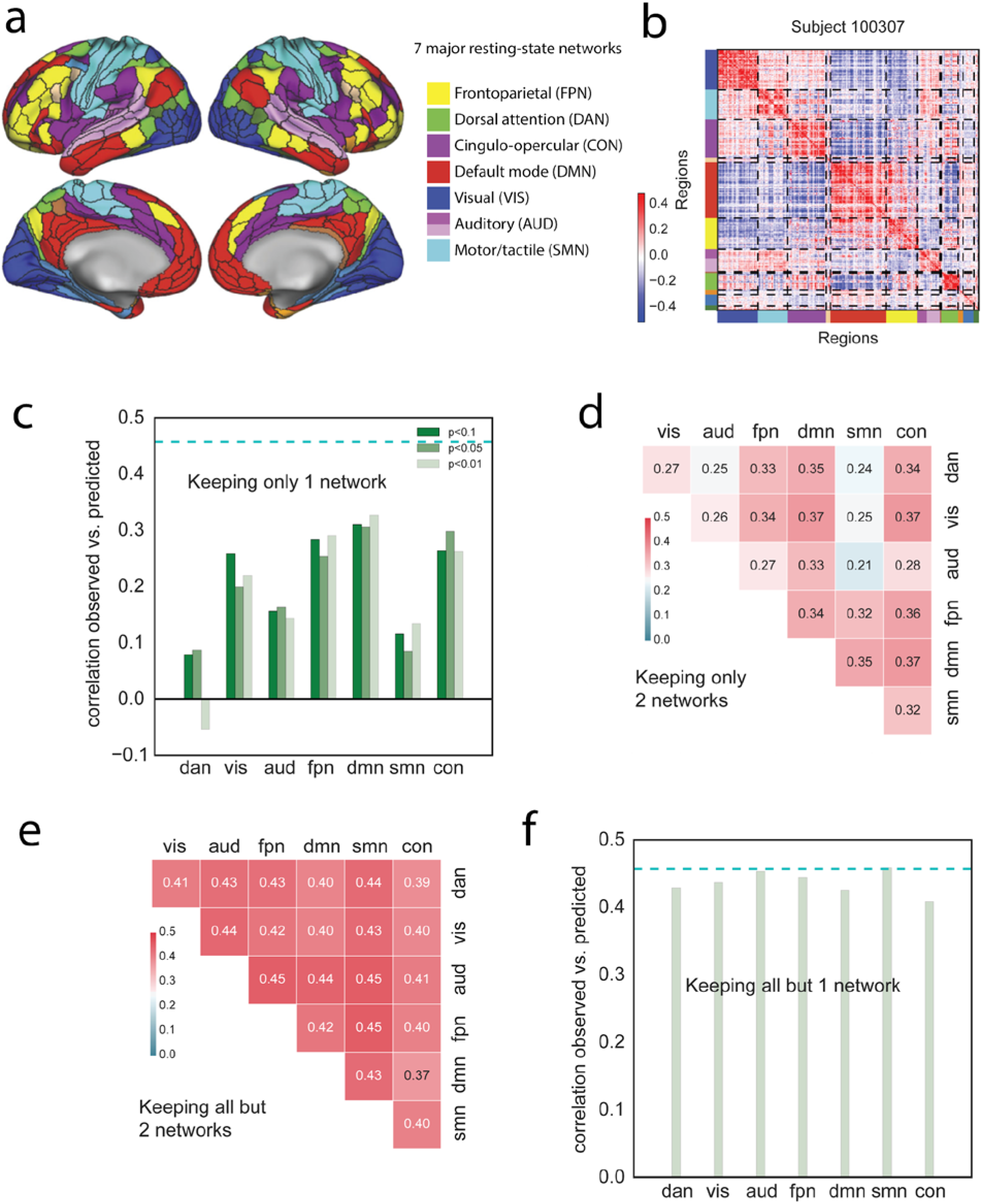
A distributed neural basis for g. **a)** Assignment of parcels to major resting-state networks, reproduced from Ito et al. [71]. **b)** Example REST12 functional connectivity matrix ordered by network, for an individual subject (id=100307). **c)** Prediction performance for REST12 matrices (as Pearson correlation between observed and predicted scores) using only within network edges, for the 7 main resting-state networks listed in **a**. DMN, FPN, CON and VIS carry the most information about *g*. The three shades of green correspond to three univariate thresholds for initial feature selection (we used p<0.01 as in the main analysis; as well as p<0.05 and p<0.1 to make sure that the results were not limited by the inclusion of too few edges). The dashed cyan line shows for comparison the prediction performance for the whole-brain connectivity matrix (same data as in **Figure 2**). **d)** Prediction performance with two networks only (univariate feature selection with p<0.01). **e)** Prediction performance for REST12 matrices after “lesioning” two networks (univariate feature selection with p<0.01). **f)** Prediction performance after lesioning one network (univariate feature selection with p<0.01).

## Discussion

Deary et al. recently wrote [91]: “the effort to understand the psychobiology of intelligence has a resemblance with digging the tunnel between England and France: We hope, with workers on both sides having a good sense of direction, that we can meet and marry brain biology and cognitive differences.” The present study is a step in this direction, offering the to-date most robust investigation specifically focused on predicting intelligence from resting-state fMRI data. Here we used factor analysis of the scores on 10 cognitive tasks to derive a bi-factor model of intelligence, including a common g-factor and broad ability factors, which is the standard in the field of intelligence research [46,92]. We used reliable estimates of functional connectivity in a large sample of subjects, from close to one hour of high-quality resting-state fMRI data per subject. We used the best available inter-subject alignment algorithm (MSM-All), a stringent control for confounding variables, and out-of-sample prediction. With these state-of-the-art methods on both ends of the tunnel, we demonstrated a strong relationship between general intelligence *g* and resting-state functional connectivity (at least as strong as the well established relationship of intelligence with brain size [44,46]).

We further established that predictive network edges were fairly distributed throughout the brain, though they mostly fell within 4 of the 7 major resting-state networks: the frontoparietal network, the default mode network, the control network, and the visual network. These findings are in general agreement with the parieto-frontal integration theory (P-FIT) of intelligence. This neuroanatomical description of informative edges should be considered preliminary, as we have yet to explore how it is affected by analytical choices, such as the brain parcellation scheme and the predictive model. Removing the nodes from one or two networks had little effect on prediction scores (**Figure 3 e,f**), supporting the conclusion that information is quite distributed. This latter finding might seem at odds with prior reports that focal lesions in frontal or parietal lobe are correlated with reduced intelligence [61]. However, there is a fundamental difference here: actual neurological lesions [93] would remove the function of the lesioned regions, and would thus remove information not only for edges directly linked to the lesioned parcels, but remove additional information in complex ways all across the brain. Our virtual “lesions” do not do this and instead retain information from the “lesioned” regions that is broadcast across other brain regions.

Our findings considerably extend a previous report [40] which had hinted at a relationship between resting-state functional connectivity and intelligence using a much smaller subject sample (N=117), no account of potential confounding variables, a cross-validation scheme that did not respect family structure, a less functionally accurate inter-subject alignment, and a lower quality measure of intelligence (the short modified version of Raven’s Progressive Matrices). We indeed found evidence that measurement quality, both on the behavioral and on the neural side, moderated the effect size of the relationship between brain and behavior [46]. We achieved better prediction performance using *g* rather than the number of correct responses on the PMAT24_A test; and we achieved better performance using REST12 matrices (~1h of data per subject) rather than REST1 or REST2 matrices (~30min of data per subject). Though this is of course expected statistically -- the noise ceiling gets lower as noise increases for the two variables that are correlated -- this is an important observation for future explorations in other datasets. In many aspects the neural and behavioral data in the HCP are of higher quality than most other large-scale neuroimaging projects; conducting similar analyses in other datasets may yield smaller effect sizes solely because of lower data quality and, just as importantly, quantity. Despite this caveat, and despite the care that we took to use crossvalidated predictions to assess out-of-sample generalizability, it will be important to replicate our finding in an independent dataset, if only to establish the bounds of the generalizability of our findings. Though beyond the scope of this study, we are already exploring three candidate publicly available datasets with suitable imaging and behavioral assessment in large cohorts of subjects: the Cambridge Center for Ageing and Neuroscience (Cam-CAN, [94]); the Nathan Kline Institute Rockland sample [95]; and the UK Biobank [96].

The general factor of intelligence *g* that we derived from 10 cognitive task scores is as reliable as it can be in this dataset, and would be unlikely to improve substantially even if additional measures were available. An interesting question for future studies will be to look at the predictability of the subscales: crystallized ability, processing speed, visuospatial reasoning, and memory. Addition of tasks in each of these subdomains would increase the reliability of these specific ability factors, and allow for a more precise exploration of their neural bases. This would require a much longer assessment and many more ability tests. Rather than build an entirely new dataset from scratch, the possibility of testing all HCP subjects again on a lengthier, diverse cognitive ability battery should be considered [97].

Another factor that can moderate the relationship between variables is range restriction of variables [98]. Here, there is some concern that the range of intelligence scores in the HCP subject sample may be restricted to the higher end of the distribution. While published normative data is currently unavailable for the Penn matrix reasoning test and other Penn CNB tests in the age range of our subjects, the NIH toolbox tests provide age-normed scores. Inspection of these scores indicates that the HCP subject sample is indeed biased towards higher scores (in particular for crystallized abilities; see **Supplementary Figure 4**). This sampling bias is a well-known, systemic issue in experimental psychology [99], and one that is difficult to avoid despite efforts to recruit from the entire population. A natural question to ask is whether the neural bases of mental retardation, and of genius at the other end of the spectrum, lie in the same continuum as what we describe in this study. For instance, it is well known that macrocephaly (unusually large brains) can also be associated with mental retardation, so that the association of intelligence with larger brain size only holds within the normal range. Future studies in samples with a larger range of intelligence should explore this important question.

Though we qualified our approach as state-of-the-art on the behavioral side, intelligence researchers would likely object that deriving factor scores is a thing of the past, and that analyses should be conducted in latent space (in part because of factor indeterminacy). This objection can mostly be ignored in our situation, where we only looked at the general factor *g*, given that it can be so precisely estimated in our dataset. However to study specific factors we should consider casting the brain-behavior relationship problem within the framework of structural equation modeling (see [30] for an example).

We were very careful to regress out several potential confounds [86], such as brain size, gender and age, before performing predictions. While this gives us some comfort that the results reported here are indeed specific to general intelligence, the confound regression approach could certainly be improved further. There are two main concerns: one the one hand, we may be throwing out relevant variance and injecting noise into the *g* scores by bluntly regressing out confounding variables -- a more careful cleanup should be attempted, for example using a well-specified structural equation model [100]. However, the approach we took is superior to ignoring the issue of potential confounds altogether (as prior studies have done), which is likely to inflate predictability and compromise interpretation. On the other hand, the list of confounding variables that we considered was not exhaustive: for example, we did not regress out variance in the Openness factor of personality, which we have previously found to be correlated with intelligence [43]; we also did not consider effects of educational achievement or socio-economic status; and there are certain to be other confounds that were not measured at all. Furthermore, it is very likely that regressing out the mean framewise displacement does not properly account for non-linear effects of motion. Cleaning resting-state fMRI data of the effects of motion remains a very intense topic of research for studies of brain-behavior relationships [101–103]. While we are confident that our current results are not solely explained by motion in the scanner, a full quantification of this issue remains warranted.

It is worth mentioning a related, entirely data-driven study that was recently conducted by Smith and colleagues on an earlier release of the same HCP dataset (N=461) [104]. Using canonical correlation analysis (CCA), the authors demonstrated that a network of brain regions that closely resembled the default mode network was highly related to a linear combination of behavioral scores that they label a “positive-negative mode of population covariation”. In essence, this combination is a neurally derived general factor, encompassing cognitive and other behavioral tasks. Our study is in general agreement with these results, as we found high predictability of general intelligence *g* from connections within the default mode network.

Where do we go from here? As we know that task functional MRI [31,34] and structural MRI data (brain size [44], as well as morphometric features [63]) also hold information that is predictive of cognitive ability, a natural question is whether combining functional and structural data will allow us to account for more variance in the general intelligence factor *g*. The more variance we can account for, the more trustworthy and thus interpretable our models become, and we can hope to further refine our understanding of the neural bases of general intelligence.

Of course, mere prediction does not yet illuminate mechanisms, and we would ultimately wish to have a much more detailed causal model that explains how genetic factors, brain structure, brain function, and individual differences in variables such as *g* and personality relate to real-life outcomes. Given that *g* is already known to predict outcomes such as lifespan and salary, a structural model incorporating all of these variables should provide us with the most comprehensive understanding of the mechanisms, and the most effective information for targeted interventions.

Finally, we would like to situate this paper in the broader context of this special issue. Intelligence can be quantified across species and is highly heritable. Are similar brain networks the most predictive of variability in intelligence across mammals? Are there measures of heritability or brain structure, as compared to brain function, that might be better predictors in some species than others? It would be intriguing to find that humans share with other species a core set of genetically specified constraints on intelligence, but that humans are unique in the extent to which education and learning can modify intelligence through the incorporation of additional variability in brain function.

## Author contributions

J. Dubois and P. Galdi developed the overall general analysis framework and conducted some of the initial analyses for the paper. J. Dubois conducted all final analyses and produced all figures. L. Paul helped with literature search, analysis of behavioral data, and interpretation of the results. J. Dubois and R. Adolphs wrote the initial manuscript and all authors contributed to the final manuscript. All authors contributed to planning and discussion on this project. The authors declare no conflict of interest.

## Acknowledgments

We thank Stuart Ritchie, Gilles Gignac, William Revelle, and Ruben Gur for invaluable advice on the behavioral side of the analyses -- though the final analytical choices rest solely with the authors.

## Data Sharing

The Young Adult HCP dataset is publicly available at https://www.humanconnectome.org/study/hcp-young-adult. Analysis scripts are available in the following public repository: https://github.com/adolphslab/HCP_MRI-behavior.

## Supplementary material

**Supplementary Table 1.** List of cognitive measures included in our analyses.

| Test Name | Variable(s) | Test Activity / Outcome |
| --- | --- | --- |
| NIH Toolbox Dimensional Change Card Sort Test | **CardSort_Unadj**<br>[Unadjusted Scale Score] | Two target pictures are presented that vary along two dimensions (e.g., shape and color). Participants are asked to match a series of bivalent test pictures (e.g., yellow balls and blue trucks) to the target pictures, first according to one dimension (e.g., color) and then, after a number of trials, according to the other dimension (e.g., shape). “Switch” trials are also employed, in which the participant must change the dimension being matched. For example, after 4 straight trials matching on shape, the participant may be asked to match on color on the next trial and then go back to shape, thus requiring the cognitive flexibility to quickly choose the correct stimulus. Scoring is based on a combination of accuracy and reaction time, and the test takes approximately 4 minutes to administer. |
| NIH Toolbox Flanker Inhibitory Control and Attention Test | **Flanker_Unadj**<br>[Unadjusted Scale Score] | The test requires the participant to focus on a given stimulus while inhibiting attention to arrows flanking it. Sometimes the middle stimulus is pointing in the same direction as the “flankers” (congruent) and sometimes in the opposite direction (incongruent). Scoring is based on a combination of accuracy and reaction time, and the test takes approximately 3 minutes to administer. |
| NIH Toolbox List Sorting Working Memory Test | **ListSort_Unadj**<br>[Unadjusted Scale Score] | This task assesses working memory and requires the participant to sequence different visually- and orally-presented stimuli. Pictures of different foods and animals are displayed with both a sound clip and written text that name the item. The task has two different conditions: 1-List and 2-List. In the 1-List condition, participants are required to order a series of objects (either food or animals) in size order from smallest to largest. In the 2-List condition, participants are presented both food and animals and are asked to report the food in size order, followed by the animals in size order. Participants have two practice items, in which the images briefly “flash” sequentially on the screen in each condition. |
| NIH Toolbox Picture Sequence Memory Test | **PicSeq_Unadj**<br>[Unadjusted Scale Score] | The Picture Sequence Memory Test is a measure developed for the assessment of episodic memory. It involves recalling increasingly lengthy series of illustrated objects and activities that are presented in a particular order on the computer screen. The participants are asked to recall the sequence of pictures that is demonstrated over two learning trials; sequence length varies from 6-18 pictures, depending on age. Participants are given credit for each adjacent pair of pictures (i.e., if pictures in locations 7 and 8 and placed in that order and adjacent to each other anywhere – such as slots 1 and 2 – one point is awarded) they correctly place, up to the maximum value for the sequence, which is one less than the sequence length (if there are 18 pictures in the sequence, the maximum score is 17, because that is the number of adjacent pairs of pictures that exist). The test takes approximately 7 minutes to administer. |
| NIH Toolbox Picture Vocabulary Test | **PicVocab_Unadj**<br>[Unadjusted Scale Score] | This measure of receptive vocabulary is administered in a computerized adaptive format. The respondent is presented with an audio recording of a word and four photographic images on the computer screen and is asked to select the picture that most closely matches the meaning of the word. This test takes approximately 4 minutes to administer. |
| NIH Toolbox Pattern Comparison Processing Speed Test | **ProcSpeed_Unadj**<br>[Unadjusted Scale Score] | This test measures speed of processing by asking participants to discern whether two side-by-side pictures are the same or not. Participants’ raw score is the number of items correct in a 90-second period. The items are designed to be simple to most purely measure processing speed. The test overall takes approximately 3 minutes to administer. |
| NIH Toolbox Oral Reading Recognition Test | **ReadEng_Unadj**<br>[Unadjusted Scale Score] | The participant is asked to read and pronounce letters and words as accurately as possible. The test administrator scores them as right or wrong. The test is given via a computerized adaptive format and requires approximately 3 minutes. |
| Penn CNB Penn Progressive Matrices | **PMAT24_A_CR**<br>[# of Correct Resp] | Fluid intelligence is measured using Raven’s Progressive Matrices . We use Form A of an abbreviated version of the Raven’s developed by Gur and colleagues. Participants are presented with patterns made up of 2x2, 3x3 or 1x5 arrangements of squares, with one of the squares missing. The participant must pick one of five response choices that best fits the missing square on the pattern. The task has 24 items and 3 bonus items, arranged in order of increasing difficulty. However, the task discontinues if the participant makes 5 incorrect responses in a row. |
| Penn CNB Penn Word Memory Test | **IWRD_TOT**<br>[Total # of Correct Resp] | Verbal episodic memory is measured using Form A of the Penn Word Memory Test. Participants are shown 20 words and asked to remember them for a subsequent memory test. They are then shown 40 words (the 20 previously presented words and 20 new words matched on memory related characteristics). They decide whether they have seen the word previously by choosing among “definitely yes,” “probably yes,” “probably no,” and “definitely no.” |
| Penn CNB Variable Short Penn Line Orientation | **VSPLOT_TC**<br>[total # correct] | Spatial orientation processing is measured using the Variable Short Penn Line Orientation Test. Participants are shown two lines with different orientations. They have to rotate one of the lines (a moveable blue one) so that is parallel to the other line (a fixed red line). The rotation of the blue line is accomplished by clicking buttons on the keyboard that rotate the lines either clockwise or counterclockwise. Across trials, the lines vary in their relative location on the screen, though the distance between the centers of the two lines is always the same. The length of the red line is always the same, but the length of the blue line can be either short or long. There are a total of 24 trials |

### Confirmatory factor analysis

Though deriving factor scores from an EFA is often done by empirical researchers, it is theoretically preferable to use a confirmatory factor analysis (CFA) framework. A major difference is that a CFA model typically restricts cross-loadings (an observed variable loading on several latent factors), while the EFA allows them; this can reduce the size of the *g* factor in the EFA.

Using the EFA solution, we specified a bi-factor model, including a general factor (loading on all tasks) and four group factors (loading on subsets of tasks), in a confirmatory factor analysis. The model does not allow for any cross-loadings of a task on several factors, and group factors are orthogonal to one another and to the general factor. As some of the group factors are only defined by two indicators, it was necessary to impose a constraint for the purposes of identification. We fixed the unstandardized loadings for both tasks to 1.0 in this case. We initially found that the lavaan model did not converge, and identified that the issue lied with the ListSort_Unadj task score. We ran the CFA without ListSort_Unadj, with the following lavaan syntax:

~~~
*#g-factor
g    =~ CardSort_Unadj + Flanker_Unadj + ProcSpeed_Unadj + PicVocab_Unadj +
   ReadEng_Unadj + PMAT24_A_CR + VSPLOT_TC + IWRD_TOT + PicSeq_Unadj
#Domain factors
spd  =~ CardSort_Unadj + Flanker_Unadj + ProcSpeed_Unadj
cry  =~ 1*PicVocab_Unadj + 1*ReadEng_Unadj
vis  =~ 1*PMAT24_A_CR + 1*VSPLOT_TC
mem  =~ 1*IWRD_TOT + 1*PicSeq_Unadj
#Domain factors are not correlated with g
g ~~ 0*spd
g ~~ 0*cry
g ~~ 0*vis
g ~~ 0*mem
#Domain factors are not correlated with one another
spd ~~ 0*cry
spd ~~ 0*vis
spd ~~ 0*mem
cry ~~ 0*vis
cry ~~ 0*mem
vis ~~ 0*mem*
~~~

This CFA model converged after 49 iterations, and the fit was very good with CFI=0.974, RMSEA=0.052, SRMR=0.032, BIC = 27820.2. The standardized solution is depicted in **Supplementary Figure 1**. The general factor was found to account for 64.0% of the variance (coefficient omega_hierarchical ω_h_), while group factors accounted for 17.2% of the variance. We derived the factor scores for *g* using the regression method; we found that the scores derived from the CFA were almost perfectly correlated with the scores derived from the EFA (**Figure 1**), r=0.99.

**Supplementary Figure 1.**
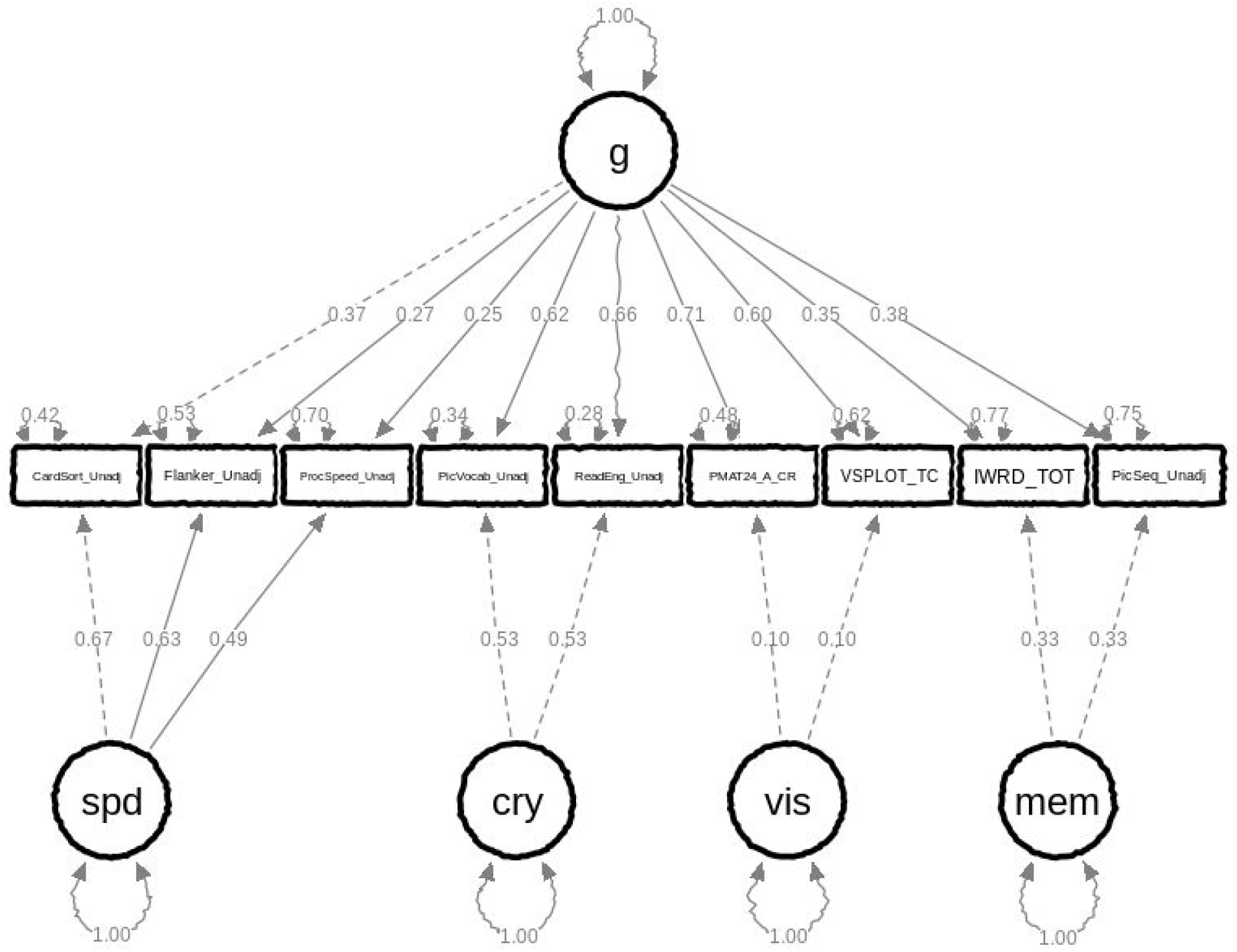
Standardized solution for our confirmatory factor analysis of the HCP cognitive task scores. The CFA omits *ListSort_Unadj* which prevented the model from converging.

**Supplementary Figure 2.**
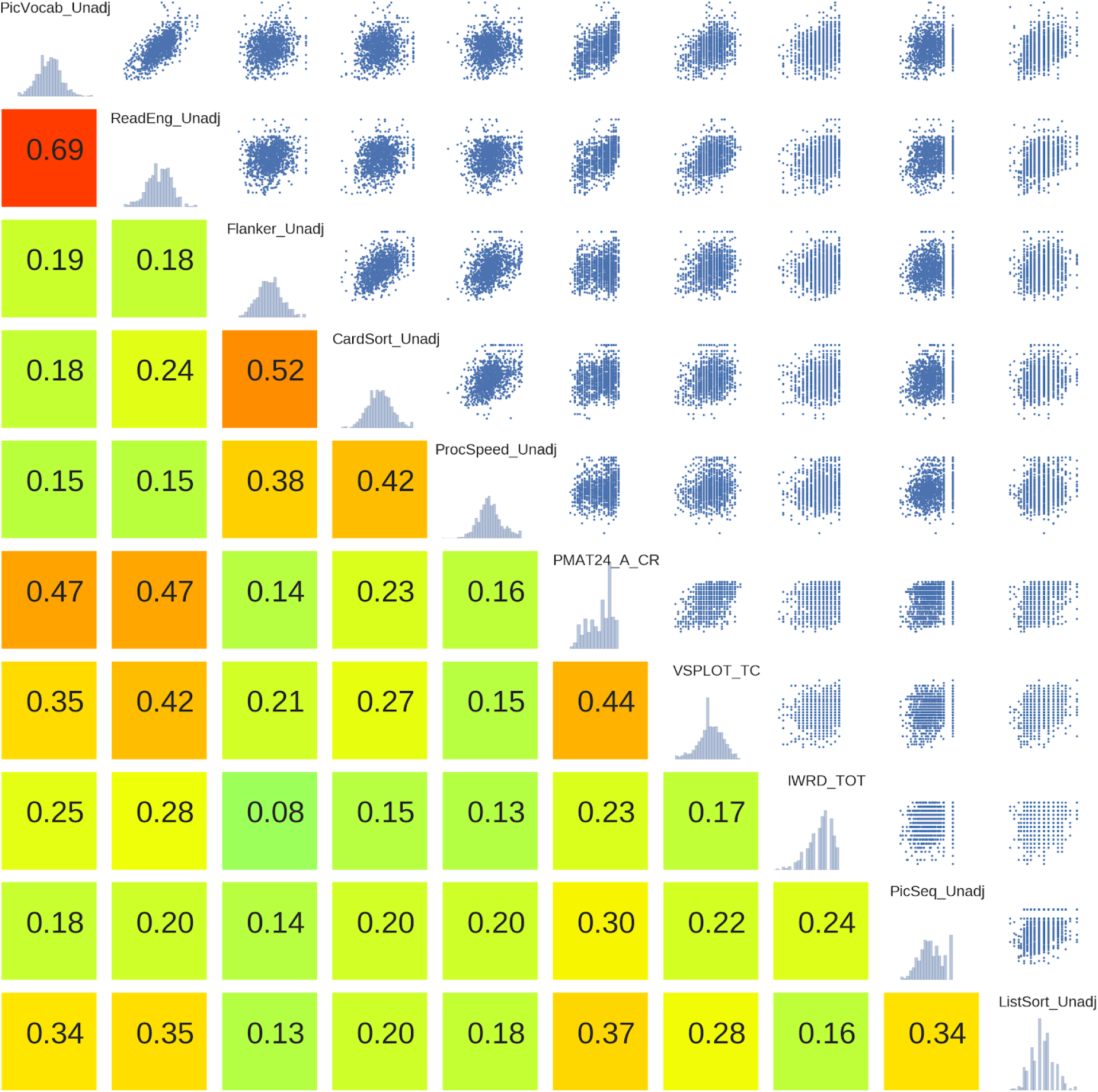
Correlations between (z-scored) HCP cognitive task scores. On the diagonal, the distribution of each of the 10 cognitive variables in shown. Below the diagonal, the Pearson correlation is displayed, together with a color visualizing the strength of the relationship. Above the diagonal, a scatter plot is displayed for each pair of variables, with x-and y-axes between −4 and 4 (standard deviation of all variables is 1 due to z-scoring).

**Supplementary Figure 3.**
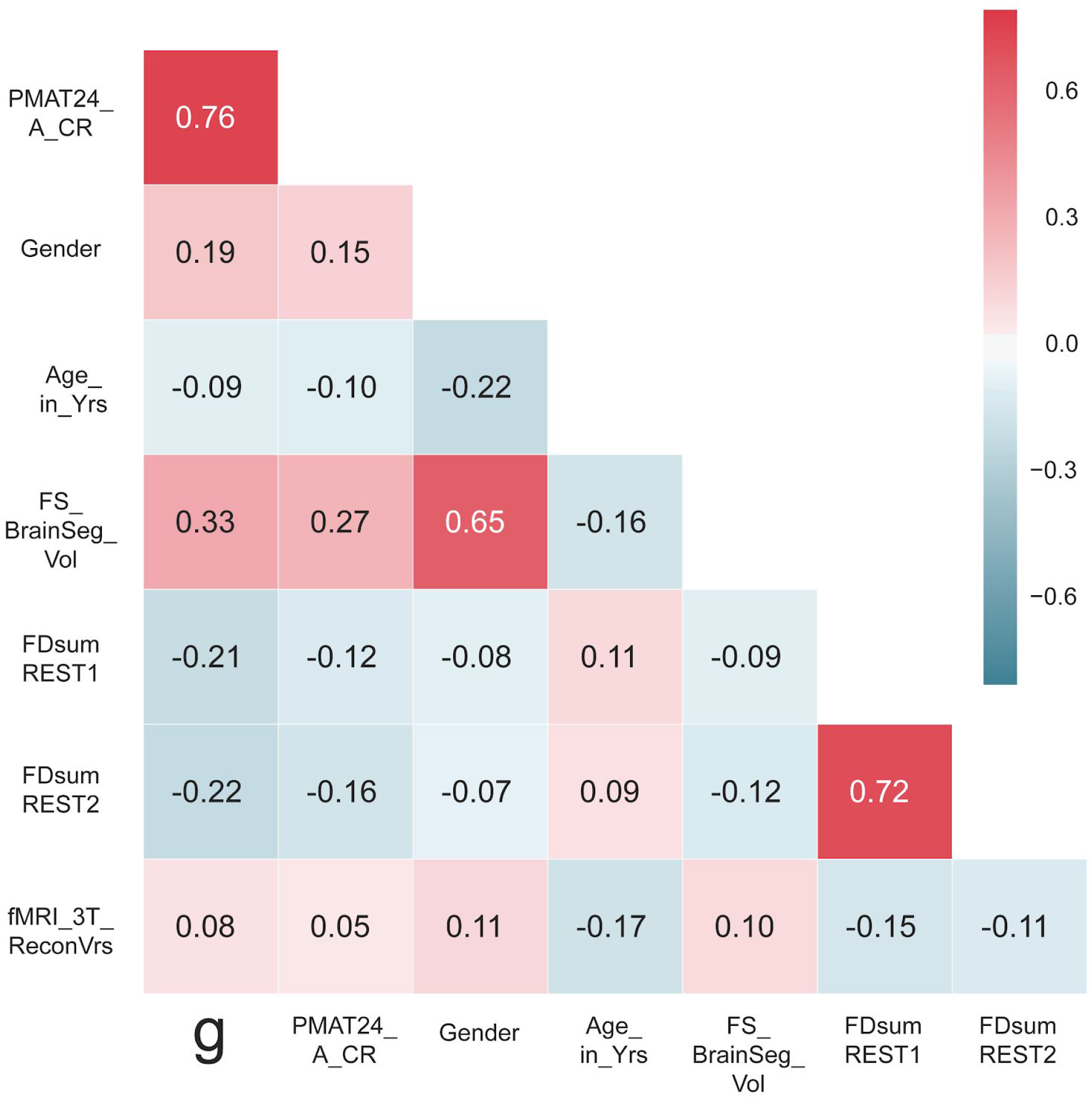
Correlation of the general intelligence factor scores with PMAT24_A_CR scores, and with potential confounds, in the sample of subjects used for prediction analyses (N=884). All of these variables except for *PMAT24_A_CR* scores were regressed out of the training set data to obtain an unconfounded measure of g.

**Supplementary Figure 4.**
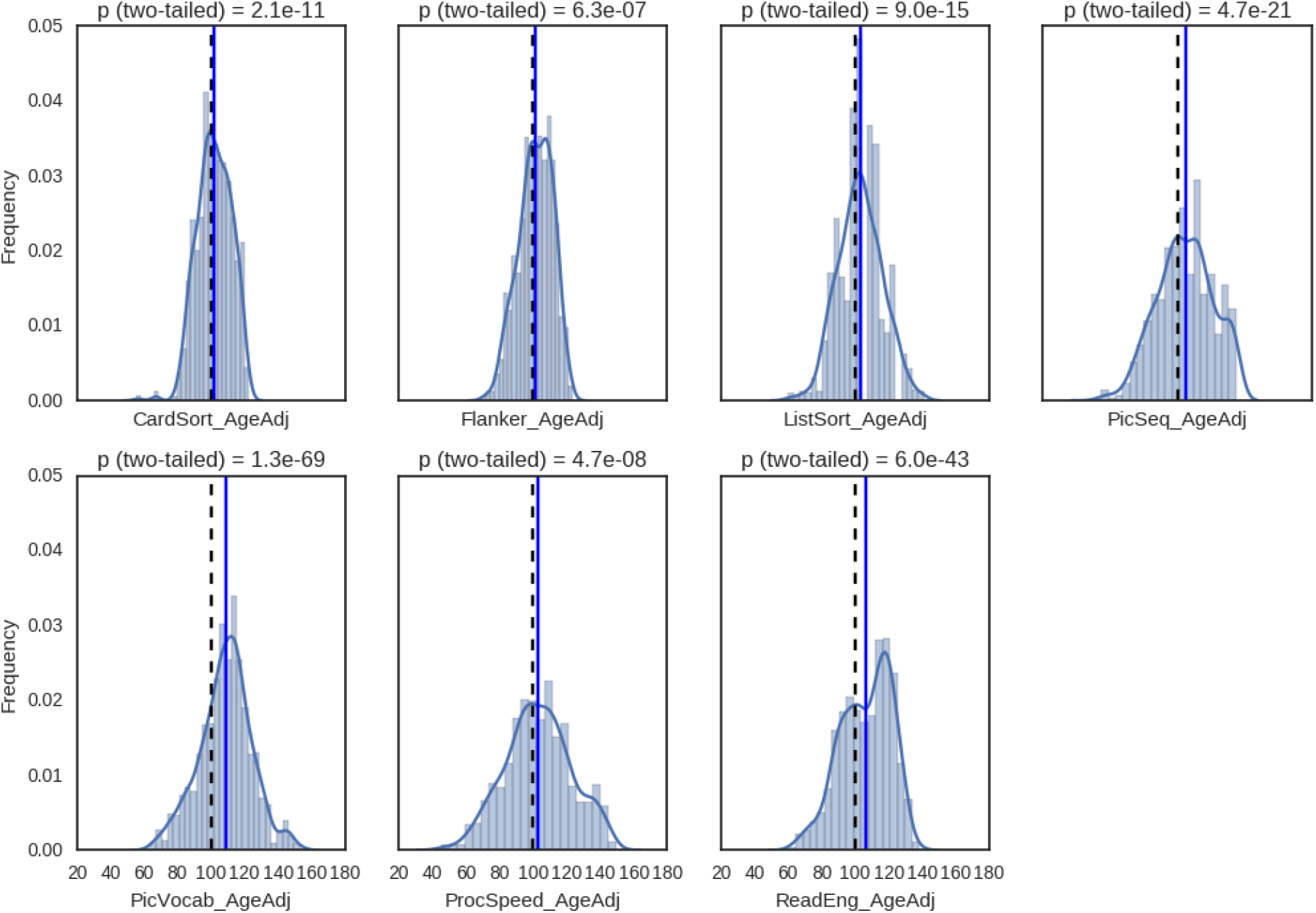
Distributions of the age-normed scores of HCP subjects on NIH-toolbox cognitive tasks. The blue line shows the mean score in our subject sample, which is greater than 100 for all tests, while the black dashed line shows the mean in the normative population. For all tests, a 1-sample Student’s t-test indicates that the mean is significantly higher than 100.

